## Supplementary Information for "Diffusion MRS tracks distinct trajectories of neuronal development in the cerebellum and thalamus of rat neonates"

[1]Wellcome Centre for Integrative Neuroimaging, FMRIB, Nuffield Department of Clinical Neurosciences, University of Oxford; [2]Cardiff University Brain Research Imaging Centre (CUBRIC), School of Psychology, Cardiff University, Cardiff, UK; [3] School of Computer Science and Informatics, Cardiff University, Cardiff, UK; [4]Mouse Imaging Centre, The Hospital for Sick Children, Toronto, Ontario, Canada; [5]Department of Medical Biophysics, University of Toronto, Toronto, Ontario, Canada

#### SUPPLEMENTARY MATERIAL

### Content

**Figure S0**. **Analytical model residuals**. Root mean square error (RMSE) computed from the results of the analytical modelling (“astrosticks and spheres”). The logarithm is displayed for visualisation: log(residuals = rmse(model fit - data)).

**Figure S1**. **Representative segmented atlases at each time point**. The reference atlas used for the longitudinal analysis is a P15-Fischer-derived-atlas. The atlas was derived from a P15-P20 registration (using a P20-Fischer-derived-atlas, itself derived from a P20-P30 registration using the original Fischer atlas). White arrows point at the thalamus and cerebellum segmentations. Red arrows point at the major registration artefacts.

**Figure S2. Cerebellar (blue) and thalamic (red) metabolic changes with age.** This figure complements Figure 3. Spectra at b=0.035 ms/$\mu m^2$ and TM=100 ms were quantified with LCModel and absolute metabolite concentrations were normalised by either the absolute voxel water signal (white columns) or the absolute voxel macromolecular concentration (grey columns).

**Figure S3. Visual comparison of diffusion properties** between regions and at each time point.

**Figure S4. Visual comparison of the parameter space** (sphere fraction, sphere radius) between the cerebellum and the thalamus at each time point for the “astrosticks + spheres” model ($D_{intra}$=0.5$\mu m^2$/ms).

**Figure S5. Parameter space (sphere fraction, sphere radius)** between the cerebellum and the thalamus at each time point for the “astrosticks + spheres” model ($D_{intra}$=0.5$\mu m^2$/ms) on averaged neonates.

**Figure S6**. **Visual regional comparison of parameters estimated from the morphometric model.** Estimations of $D_{intra}$, $N_{branch}$, $L_{segment}$ are shown for the 6 metabolites in 2 regions.

**Figure S7**. Estimations from the morphometric model of $D_{intra}$, $N_{branch}$, $L_{segment}$ are shown for the 6 metabolites in 2 regions.

**Figure S8.** DTI results in the cerebellum and thalamus. (A) Representative FA maps at P5, P15 and P30 in both regions. Images are not at scale. The image quality is particularly poor at P30, where the ghosting artefact in the ventral part of the brain appears in almost all images (poor FOV calibration). (B) Mean MD and FA evolution with age in both regions. The MD and FA values were averaged over the dMRS voxel projected onto the maps. Values come from 12 animals (6 for each region).
Given the poor image quality, we refrain from drawing conclusions from these results, but we include them in the Supplementary Information for thoroughness.
*Methods*: SE-EPI sequence (4 segments, TR=2.5 s, TE=23.5 ms, resolution: 200µm isotropic, 30 directions, b=1000 s/$mm^2$, 3 b=0 images, ∂=2.5 ms, Δ=10 ms).

**Figure S9. Parameter space (sphere fraction, sphere radius)** between the cerebellum and the thalamus at each time point for the “astrosticks + spheres” model **($D_{intra}$=0.7$\mu m^2$/ms)**

**Figure S10. Average macromolecule signal attenuation in both regions at each time point coming from the LCModel fit.** The macromolecule signal attenuation is abnormally strong in the cerebellum at P5 (black line, left). This underlines a residual motion artefact that could not be corrected during processing. The shaded error bars represent the standard deviation from the mean.

**Figure S11.** Literature estimate of $f_{sphereL,T}$ (as described in Table S2, pale rounds on the graph) as a function of age compared to $f_{sphere}$(tCr) (black triangles) $f_{sphere}$(Tau, tCho, Ins) (stars). P35 represents any age > P30. Legend: N2018 = Nakayama 2018[1]; M2019 = Miterko 2019[2]; Y2004 = Yamanaka 2004[3]; vdH2021 = van der Heijen 2021[4]; A2019 = Araujo 2019[5]; KS2014 = Kim and Scott 2014[6]. ”WM” indicates that the estimate included WM.

**Table S1:** mean relative CRLB (Cramér Rao Lower Bound) for tNAA, Glu, tCr, Tau, tCho and Ins for all diffusion conditions in both regions.

**Table S2:** estimation of $f_{sphere,LT}$ derived using the ratio of the relative thicknesses of the EGL, IGL, PL (“cell body-like” layers) and ML (“process-like” layer). “with WM” means that the white matter layer was included in the “process-like” layers (in addition to ML). Data come from manual measurements of literature cerebellar figures. References are at the bottom of the Supplementary Information.

**Supplementary information to Figure 6**

**Supplementary References**

**Figure S0**. **Analytical model residuals**. Root mean square error (RMSE) computed from the results of the analytical modelling (“astrosticks and spheres”). The logarithm is displayed for visualisation: log(residuals = rmse(model fit - data)).

Residuals: probing $D_{intra}$ using the “astrosticks” + spheres model with $D_{intra}$ fixed. $D_{intra}$ values probed = 0.3, 0.4, 0.5, 0.6, 0.7, 0.8 µm$^2$/ms. The mean value over all pups is displayed at each time point in each region (neonates fitted independently).

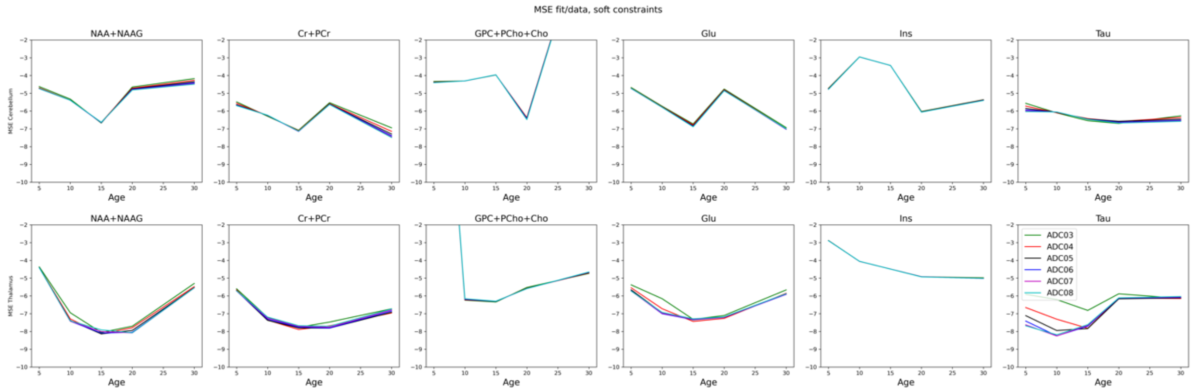

Residuals: RMSE computed for the 4 analytical models assessed: “astrosticks” (DL=1), “astrocylinders” (DL=2), “astrosticks + spheres” (DL=3) , “astrosticks + spheres” – fixed $D_{intra}$. The mean RMSE value over all pups is displayed at each time point in each region (neonates fitted independently).

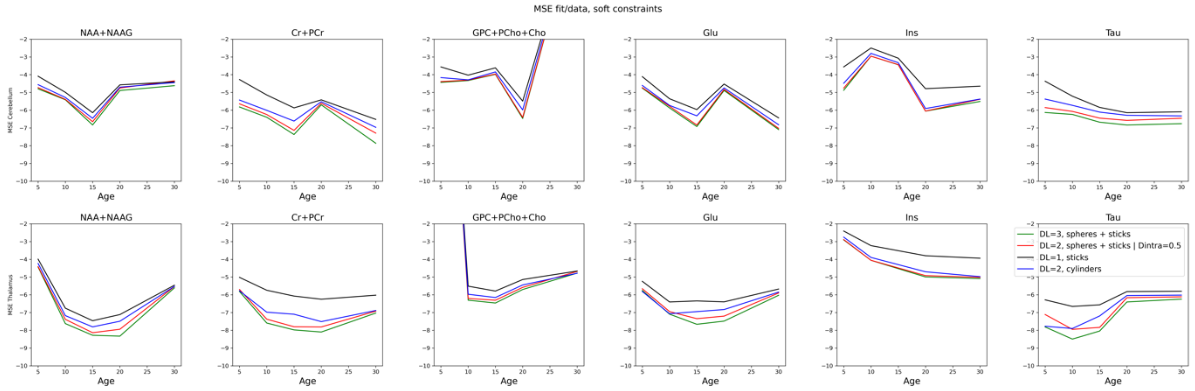

Residuals: RMSE computed from the averaged data at each time point in each region for the 4 analytical models assessed: “astrosticks” (DL=1), “astrocylinders” (DL=2), “astrosticks + spheres” (DL=3) , “astrosticks + spheres” – fixed $D_{intra}$ (neonates pooled by age & region and average fitted).

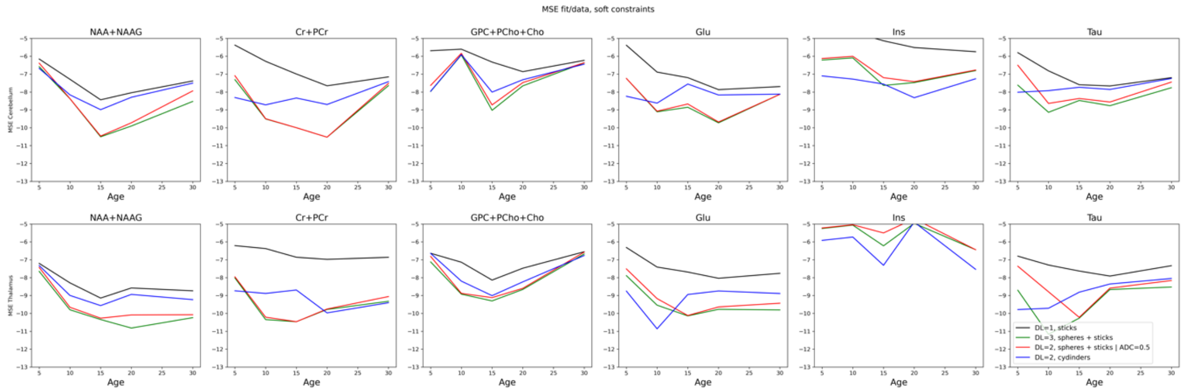

**Figure S1**. **Representative segmented atlases at each time point**. The reference atlas used for the longitudinal analysis is a P15-Fischer-derived-atlas. The atlas was derived from a P15-P20 registration (using a P20-Fischer-derived-atlas, itself derived from a P20-P30 registration using the original Fischer atlas). White arrows point at the thalamus and cerebellum segmentations. Red arrows point at the major registration artefacts.

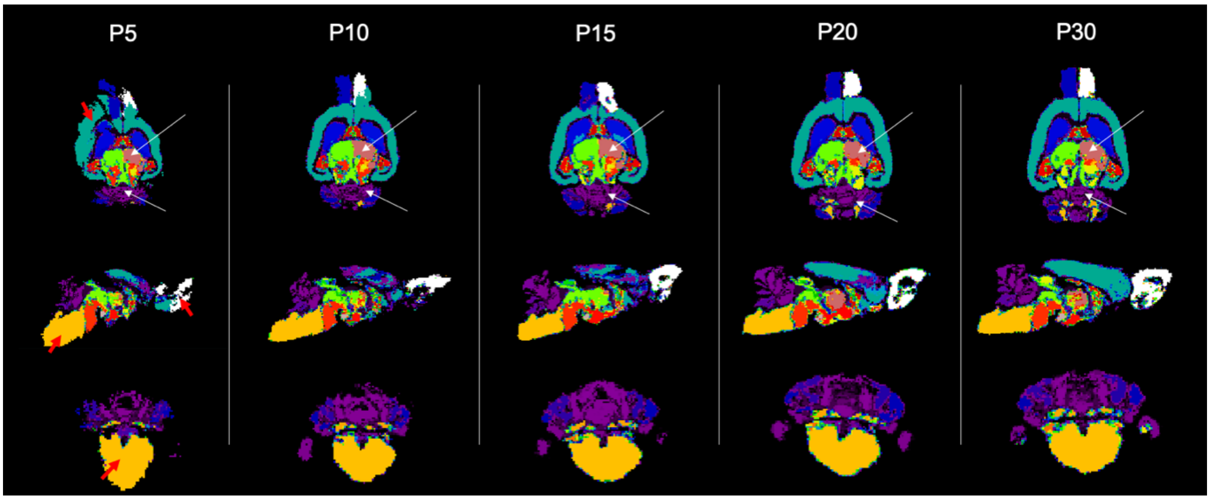

**Figure S2. Cerebellar (blue) and thalamic (red) metabolic changes with age.** This figure complements **Figure 3**.

**A.** Spectra at b=0.035 ms/µm$^2$ and TM=100 ms were quantified with LCModel and absolute metabolite concentrations were normalised by either the absolute voxel water signal (white columns) or the absolute voxel macromolecular concentration (grey columns).

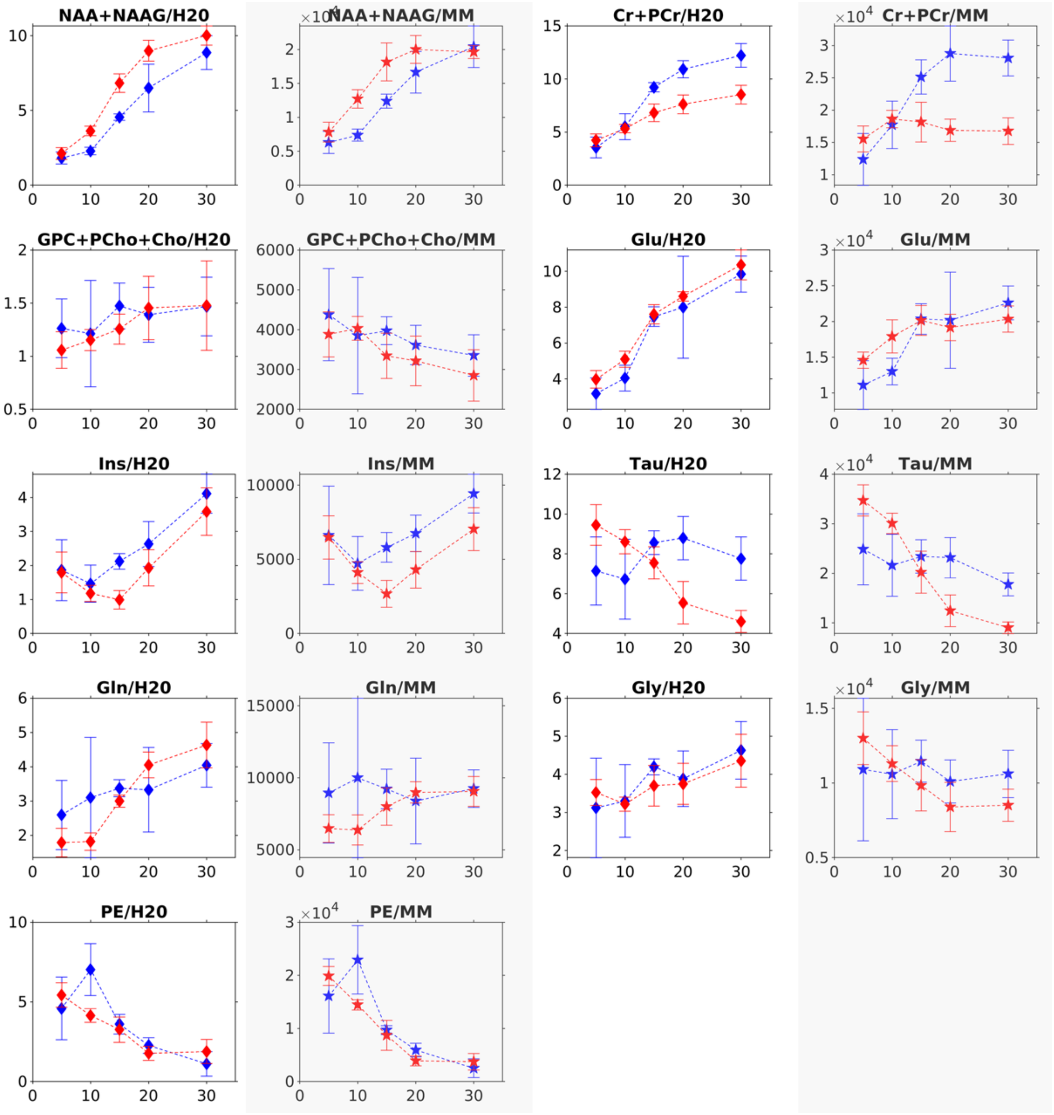

**B.** Absolute MM content in the voxel normalised by voxel size (left), raw water content normalised by voxel size (middle), and MM content normalised by water content (right). All in arbitrary units.

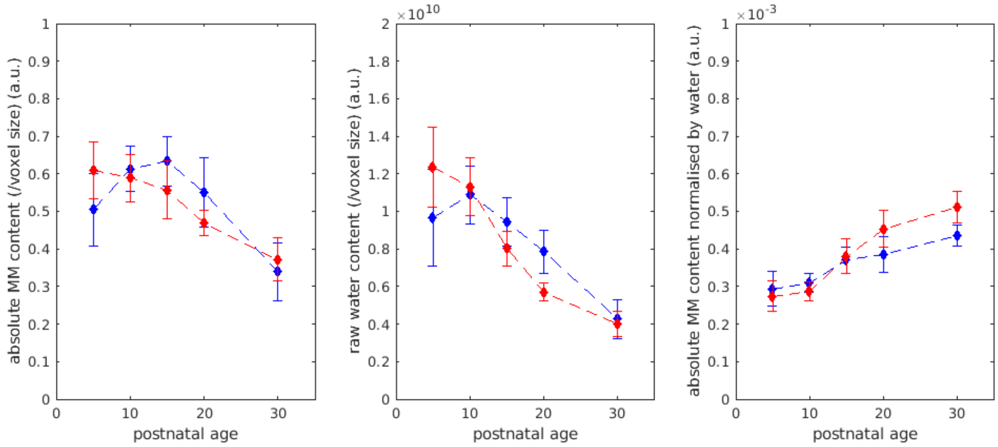

**Figure S3. Visual comparison of diffusion properties** between regions and at each time point.

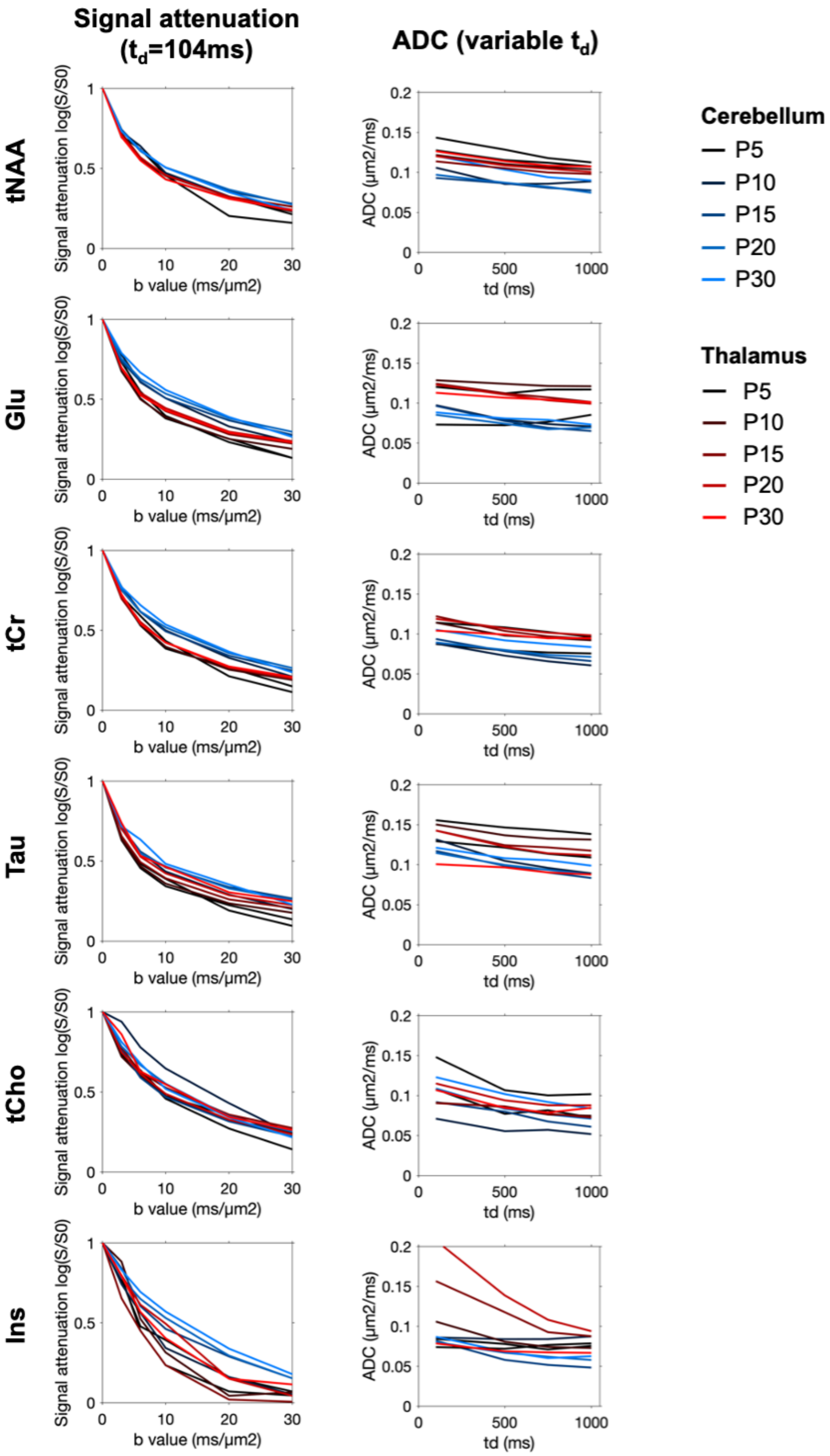

**Figure S4. Visual comparison of the parameter space** (sphere fraction, sphere radius) between the cerebellum and the thalamus at each time point for the "astrosticks + spheres" model ($D_{intra}$=0.5µm$^2$/ms).

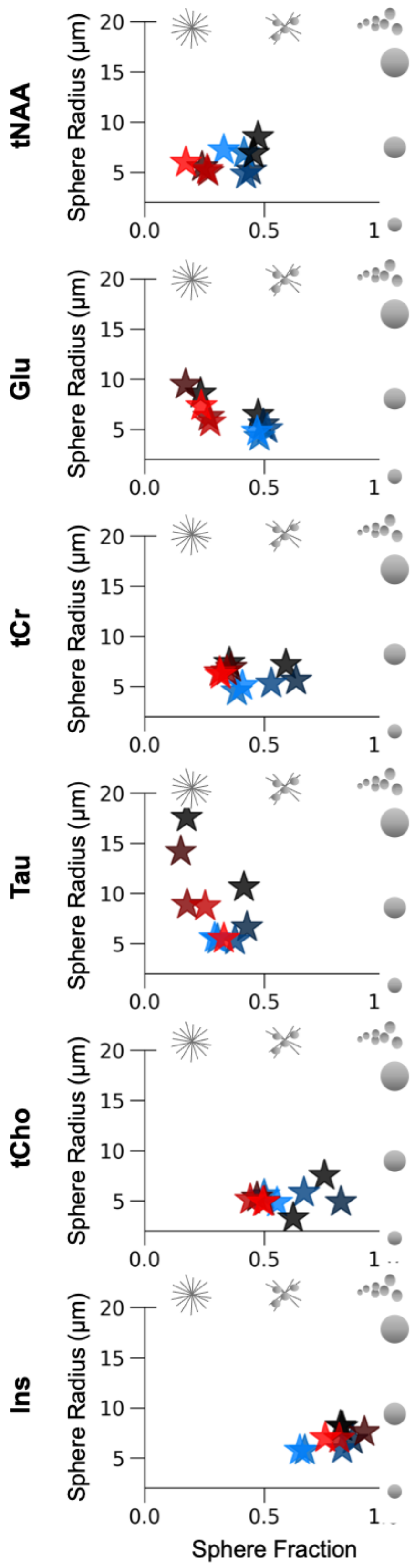

**Figure S5. Parameter space (sphere fraction, sphere radius)** between the cerebellum and the thalamus at each time point for the "astrosticks + spheres" model ($D_{intra}$=0.5µm$^2$/ms) on averaged neonates.

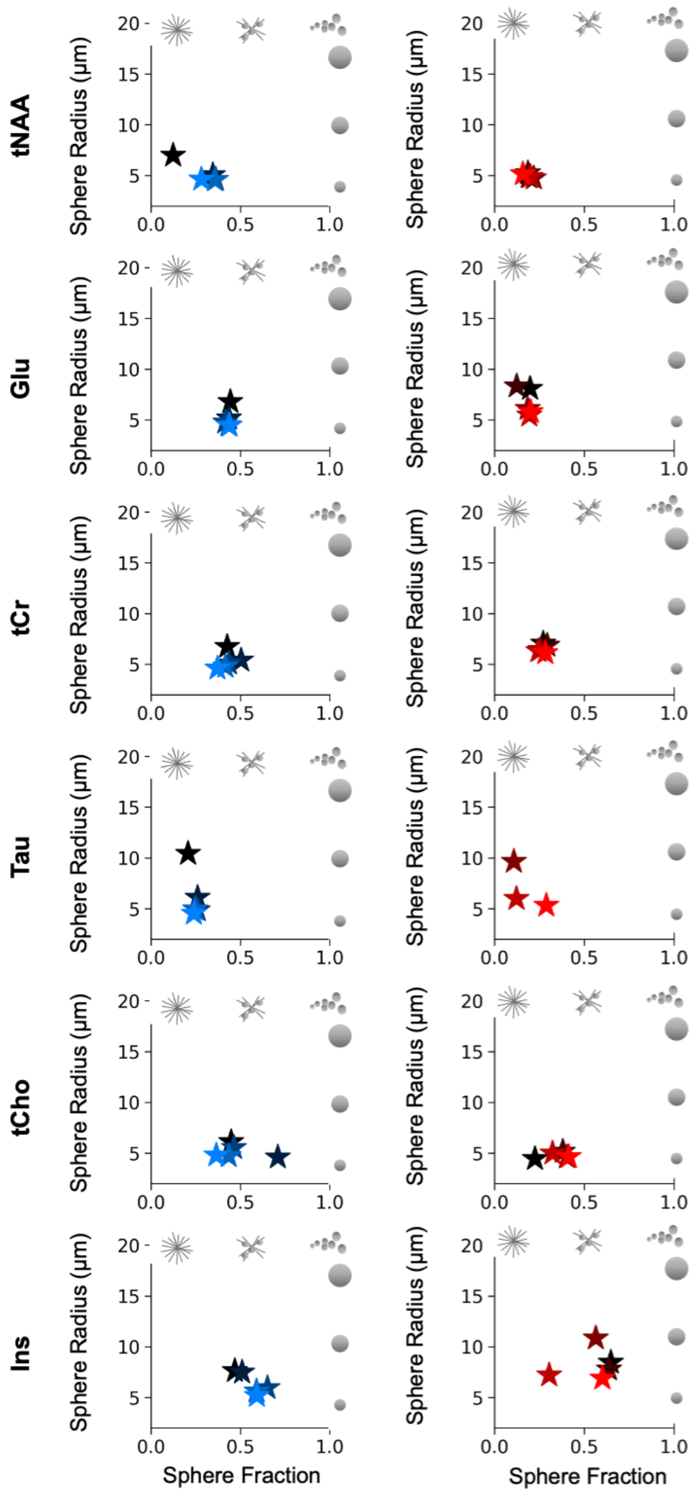

**Figure S6**. **Visual regional comparison of parameters estimated from the morphometric model.** Estimations of $D_{intra}$, $N_{branch}$, $L_{segment}$ are shown for the 6 metabolites in 2 regions.

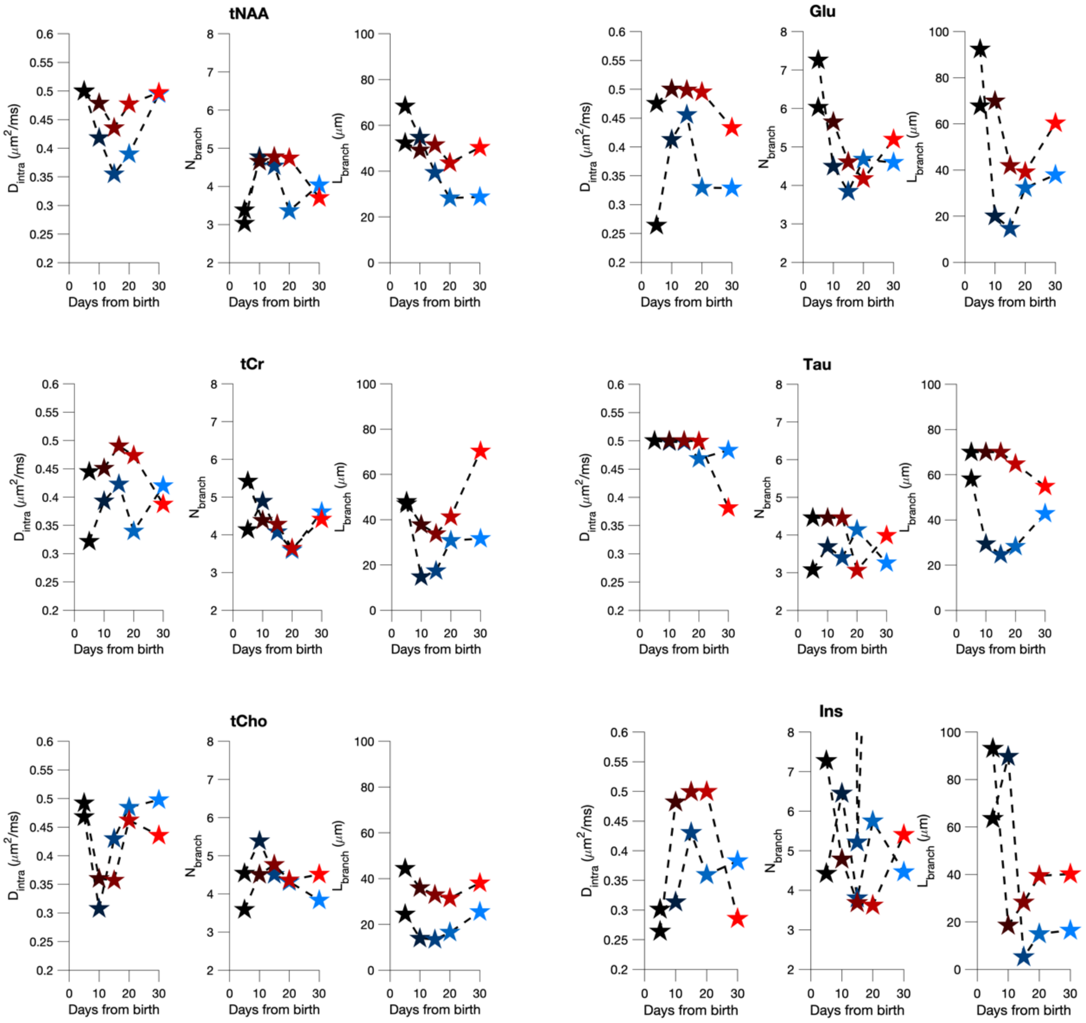

**Figure S7**. Estimations from the morphometric model of $D_{intra}$, $N_{branch}$, $L_{segment}$ are shown for the 6 metabolites in 2 regions.

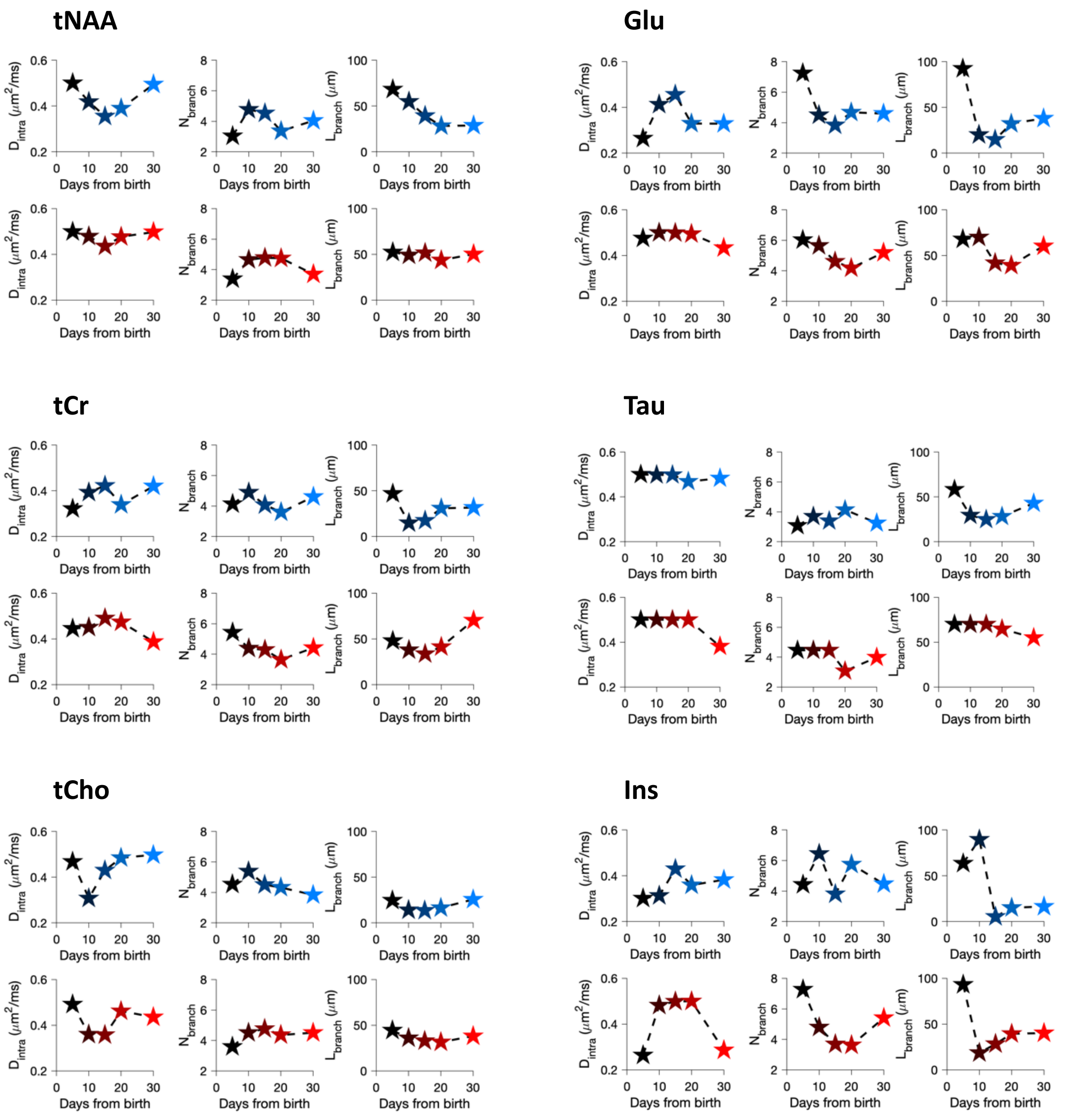

**Figure S8.** DTI results in the cerebellum and thalamus. (A) Representative FA maps at P5, P15 and P30 in both regions. Images are not at scale. The image quality is particularly poor at P30, where the ghosting artefact in the ventral part of the brain appears in almost all images. (B) Mean MD and FA evolution with age in both regions. The MD and FA values were averaged over the dMRS voxel projected onto the maps. Values come from 12 animals (6 for each region).

Given the poor image quality, we refrain from drawing conclusions from these results, but we include them in the Supplementary Information for thoroughness.

*Methods*: SE-EPI sequence (4 segments, TR=2.5 s, TE=23.5 ms, resolution: 200µm isotropic, 30 directions, b=1000 s/mm$^2$, 3 b=0 images, ∂=2.5 ms, Δ=10 ms).

**A.** FA maps from individual animals in both regions at different ages.

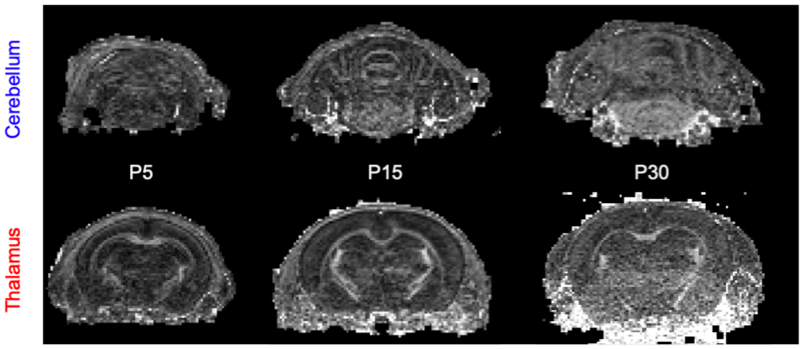

**B.** Mean MD and FA evolution with age in both regions (ROI = dMRS voxel)

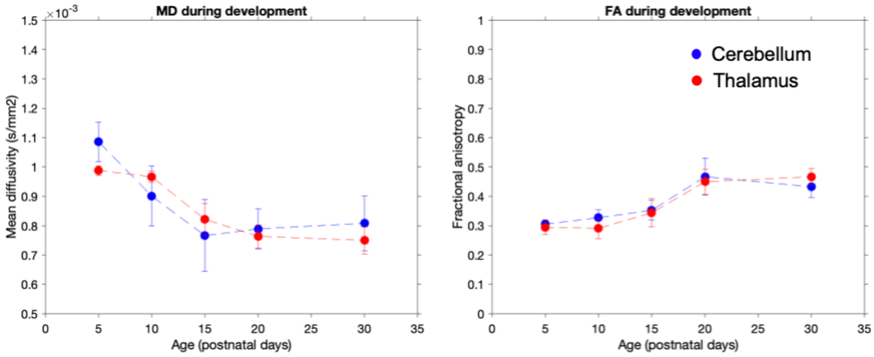

**Figure S9. Parameter space (sphere fraction, sphere radius)** between the cerebellum and the thalamus at each time point for the “astrosticks + spheres” model **($D_{intra}$=0.7µm²/ms)**

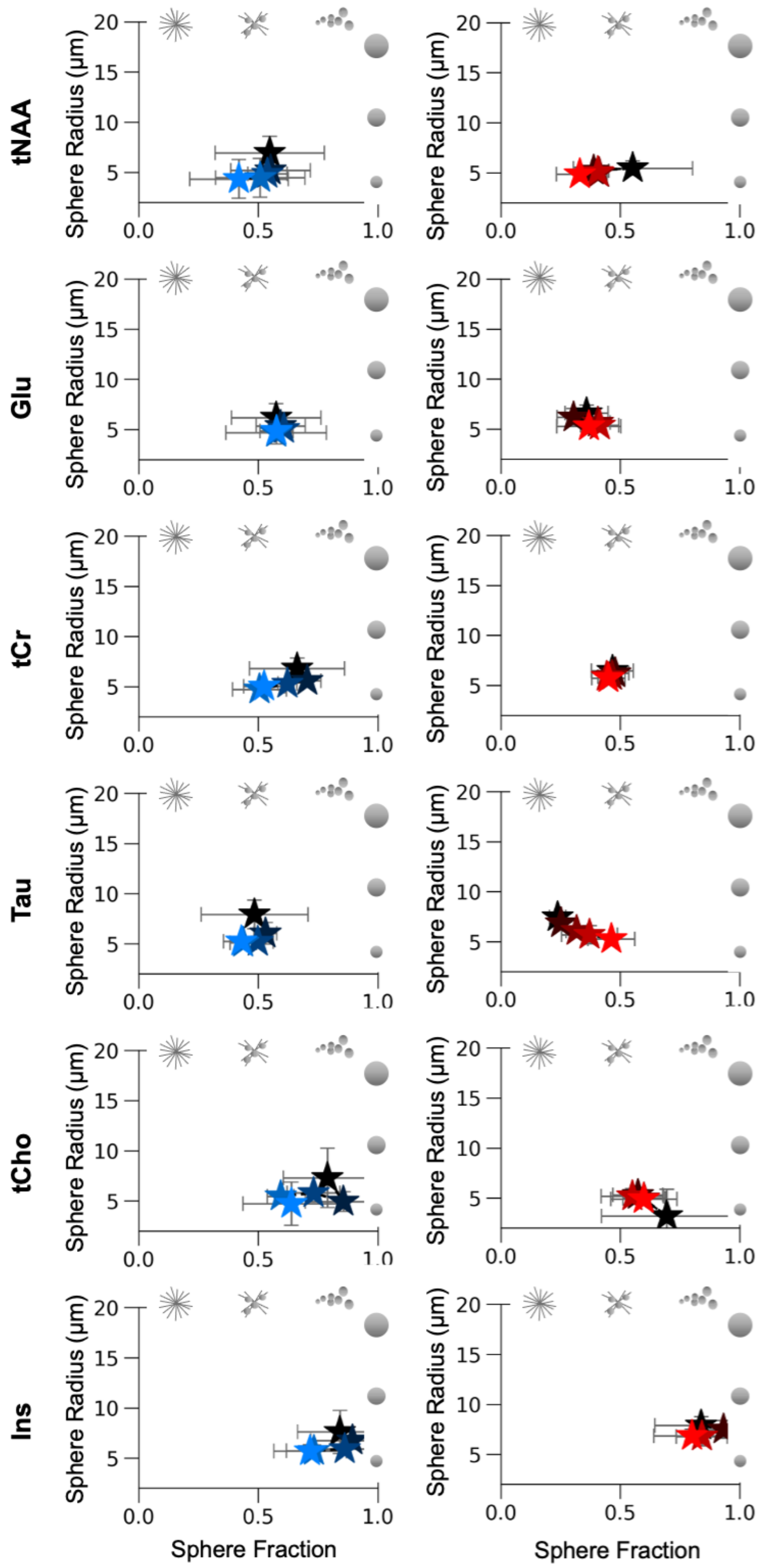

**Figure S10. Average macromolecule signal attenuation in both regions at each time point coming from the LCModel fit.** The macromolecule signal attenuation is abnormally strong in the cerebellum at P5 (black line, left). This underlines a residual motion artefact that could not be corrected during processing. The shaded error bars represent the standard deviation from the mean.

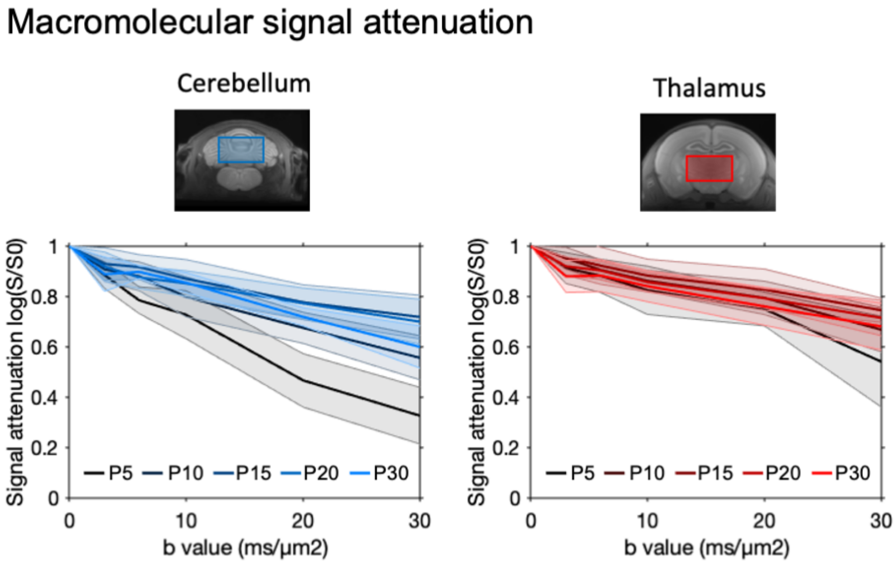

**Figure S11. Literature estimate of $f_{sphereL,T}$** (as described in Table S2, pale rounds on the graph) **as a function of age compared to $f_{sphere}$(tCr)** (black triangles) **$f_{sphere}$(Tau, tCho, Ins)** (stars)**.** P35 represents any age > P30. Legend: N2018 = Nakayama 2018[1]; M2019 = Miterko 2019[2]; Y2004 = Yamanaka 2004[3]; vdH2021 = van der Heijen 2021[4]; A2019 = Araujo 2019[5]; KS2014 = Kim and Scott 2014[6]. "WM" indicates that the estimate included WM.

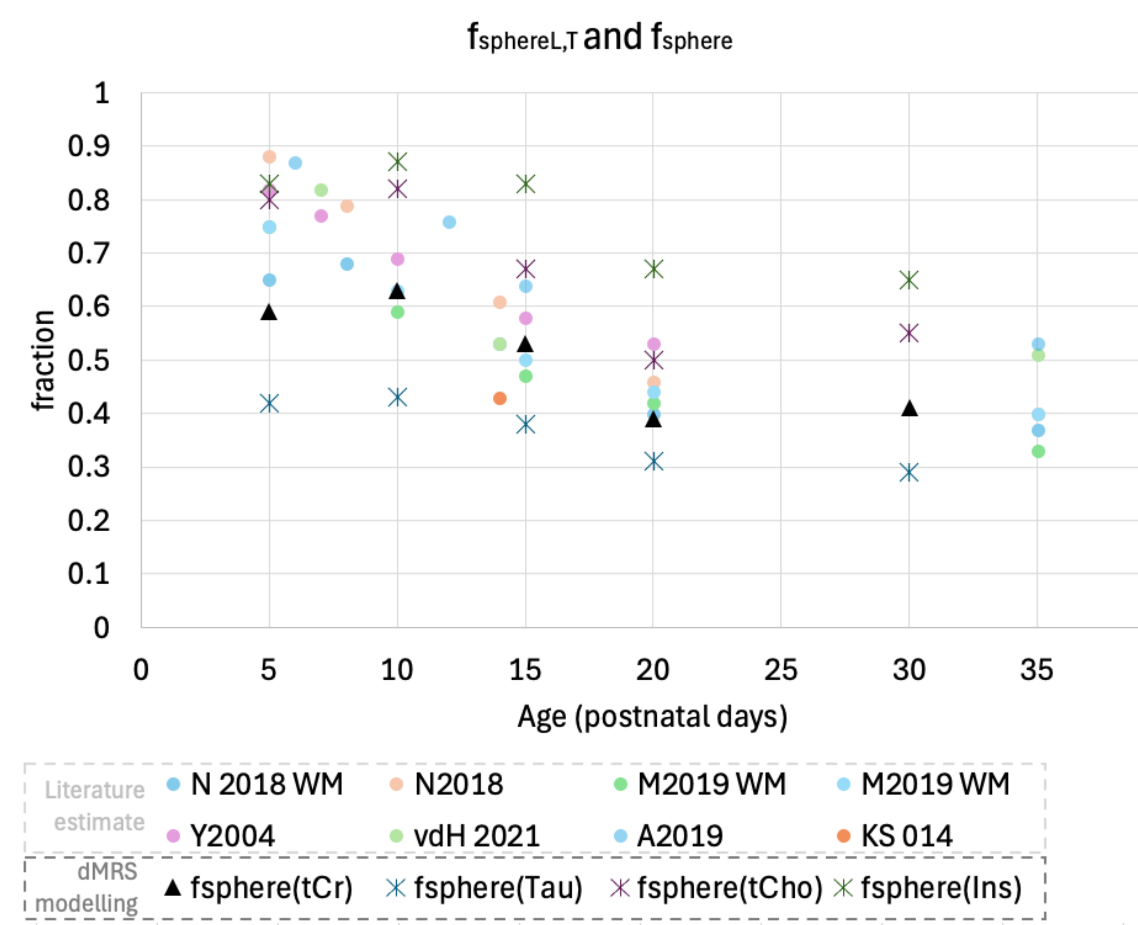

**Table S1:** mean relative CRLB (Cramér Rao Lower Bound) for tNAA, Glu, tCr, Tau, tCho and Ins for all diffusion conditions in both regions.

*HIGH B VALUES*

**CEREBELLUM**

tNAA

| b-value | P5 | P10 | P15 | P20 | P30 |
|---|---|---|---|---|---|
| 0.035 | 8.17 | 6.67 | 2.67 | 2.00 | 2.25 |
| 3.035 | 11.33 | 7.67 | 3.11 | 2.38 | 2.88 |
| 6 | 10.83 | 7.56 | 3.44 | 2.50 | 3.00 |
| 10 | 12.40 | 9.33 | 4.00 | 3.00 | 3.38 |
| 20 | 22.50 | 11.78 | 4.56 | 3.13 | 4.00 |
| 30 | 31.00 | 14.11 | 5.78 | 4.25 | 5.75 |

Glu

| b-value | P5 | P10 | P15 | P20 | P30 |
|---|---|---|---|---|---|
| 0.035 | 8.00 | 6.22 | 3.00 | 2.63 | 3.25 |
| 3.035 | 10.00 | 7.11 | 3.33 | 3.00 | 3.88 |
| 6 | 11.17 | 7.44 | 3.44 | 3.13 | 4.13 |
| 10 | 14.40 | 8.33 | 4.00 | 3.38 | 4.38 |
| 20 | 21.75 | 11.56 | 4.56 | 4.00 | 5.50 |
| 30 | 33.75 | 15.00 | 5.89 | 5.00 | 7.63 |

tCr

| b-value | P5 | P10 | P15 | P20 | P30 |
|---|---|---|---|---|---|
| 0.035 | 4.00 | 2.89 | 1.22 | 1.25 | 2.00 |
| 3.035 | 5.50 | 3.22 | 2.00 | 1.88 | 2.00 |
| 6 | 5.17 | 3.33 | 2.11 | 1.88 | 2.13 |
| 10 | 6.40 | 3.78 | 2.22 | 2.00 | 2.38 |
| 20 | 9.75 | 5.22 | 2.22 | 2.13 | 3.00 |
| 30 | 18.00 | 7.22 | 3.33 | 2.75 | 4.13 |

Tau

| b-value | P5 | P10 | P15 | P20 | P30 |
|---|---|---|---|---|---|
| 0.035 | 3.17 | 3.67 | 2.11 | 2.25 | 3.38 |
| 3.035 | 4.17 | 4.22 | 2.56 | 2.75 | 4.13 |
| 6 | 4.33 | 4.44 | 2.67 | 2.75 | 4.00 |
| 10 | 5.00 | 5.11 | 3.33 | 3.13 | 5.00 |
| 20 | 8.00 | 6.56 | 3.44 | 3.63 | 5.88 |
| 30 | 27.00 | 8.67 | 4.33 | 4.50 | 8.75 |

tCho

| b-value | P5 | P10 | P15 | P20 | P30 |
|---|---|---|---|---|---|
| 0.035 | 5.83 | 7.00 | 3.67 | 4.50 | 5.00 |
| 3.035 | 7.50 | 6.11 | 3.89 | 4.75 | 5.63 |
| 6 | 7.33 | 6.00 | 4.67 | 4.88 | 5.38 |
| 10 | 8.80 | 6.89 | 5.33 | 5.13 | 6.63 |
| 20 | 9.75 | 8.67 | 6.33 | 6.50 | 8.00 |
| 30 | 20.50 | 14.33 | 8.44 | 8.25 | 12.38 |

Ins

| b-value | P5 | P10 | P15 | P20 | P30 |
|---|---|---|---|---|---|
| 0.035 | 10.50 | 15.56 | 7.00 | 6.00 | 6.38 |
| 3.035 | 12.83 | 18.11 | 7.56 | 7.00 | 7.00 |
| 6 | 20.50 | 20.89 | 9.44 | 7.00 | 7.25 |
| 10 | 22.40 | 137.56 | 12.00 | 8.63 | 8.50 |
| 20 | 267.75 | 309.44 | 17.33 | 13.50 | 12.25 |
| 30 | 533.50 | 505.89 | 133.78 | 32.75 | 21.75 |

**THALAMUS**

NAA

| b-value | P5 | P10 | P15 | P20 | P30 |
|---|---|---|---|---|---|
| 0.035 | 6.38 | 3.00 | 2.00 | 2.00 | 2.00 |
| 3.035 | 7.88 | 4.11 | 2.22 | 2.11 | 2.67 |
| 6 | 7.38 | 4.22 | 2.22 | 2.11 | 2.89 |
| 10 | 9.63 | 5.11 | 3.11 | 3.00 | 3.56 |
| 20 | 10.57 | 6.11 | 3.44 | 3.22 | 4.11 |
| 30 | 16.86 | 7.78 | 4.22 | 4.22 | 5.33 |

Glu

| b-value | P5 | P10 | P15 | P20 | P30 |
|---|---|---|---|---|---|
| 0.035 | 5.88 | 3.89 | 3.00 | 3.00 | 3.11 |
| 3.035 | 7.88 | 5.44 | 3.56 | 3.67 | 4.33 |
| 6 | 8.38 | 6.00 | 3.67 | 3.89 | 4.56 |
| 10 | 10.88 | 7.22 | 4.33 | 4.78 | 5.56 |
| 20 | 13.29 | 9.67 | 5.78 | 5.89 | 6.89 |
| 30 | 29.71 | 13.00 | 7.22 | 7.00 | 8.78 |

tCr

| b-value | P5 | P10 | P15 | P20 | P30 |
|---|---|---|---|---|---|
| 0.035 | 3.25 | 2.00 | 2.00 | 2.00 | 2.33 |
| 3.035 | 4.00 | 2.89 | 2.11 | 2.11 | 2.89 |
| 6 | 4.25 | 3.00 | 2.11 | 2.44 | 3.11 |
| 10 | 5.50 | 3.78 | 3.11 | 3.22 | 3.67 |
| 20 | 6.29 | 4.78 | 3.56 | 4.33 | 4.78 |
| 30 | 10.57 | 6.33 | 4.56 | 5.33 | 6.56 |

Tau

| b-value | P5 | P10 | P15 | P20 | P30 |
|---|---|---|---|---|---|
| 0.035 | 2.38 | 2.00 | 2.11 | 3.56 | 5.00 |
| 3.035 | 3.13 | 2.89 | 3.00 | 4.44 | 6.56 |
| 6 | 3.25 | 3.00 | 3.11 | 4.67 | 7.67 |
| 10 | 4.13 | 3.78 | 3.78 | 5.78 | 8.33 |
| 20 | 4.71 | 4.78 | 4.67 | 7.33 | 10.44 |
| 30 | 8.14 | 5.89 | 5.89 | 8.78 | 13.11 |

tCho

| b-value | P5 | P10 | P15 | P20 | P30 |
|---|---|---|---|---|---|
| 0.035 | 5.75 | 3.78 | 3.89 | 4.00 | 5.22 |
| 3.035 | 6.88 | 5.11 | 4.33 | 4.89 | 5.56 |
| 6 | 6.38 | 4.89 | 4.44 | 4.78 | 6.89 |
| 10 | 8.00 | 6.11 | 5.11 | 6.00 | 6.67 |
| 20 | 8.43 | 7.22 | 5.67 | 6.33 | 7.67 |
| 30 | 12.14 | 8.89 | 6.44 | 7.22 | 9.56 |

Ins

| b-value | P5 | P10 | P15 | P20 | P30 |
|---|---|---|---|---|---|
| 0.035 | 10.88 | 13.11 | 15.67 | 9.56 | 6.67 |
| 3.035 | 12.63 | 14.00 | 23.44 | 11.56 | 8.56 |
| 6 | 19.13 | 19.78 | 28.00 | 12.33 | 9.56 |
| 10 | 165.50 | 135.11 | 158.67 | 15.56 | 15.33 |
| 20 | 339.14 | 607.78 | 863.33 | 248.00 | 38.33 |
| 30 | 513.00 | 483.89 | 908.00 | 583.11 | 47.00 |

*LONG DIFFUSION TIMES*

**CEREBELLUM**

tNAA

| TM (b0) | P5 | P10 | P15 | P20 | P30 |
|---|---|---|---|---|---|
| 100 | 8.17 | 6.13 | 3.00 | 2.43 | 2.17 |
| 500 | 8.25 | 5.94 | 2.67 | 2.36 | 2.00 |
| 750 | 8.00 | 5.88 | 2.67 | 2.23 | 2.00 |
| 1000 | 8.57 | 6.16 | 2.75 | 2.53 | 2.04 |

| TM (b0+3) | P5 | P10 | P15 | P20 | P30 |
|---|---|---|---|---|---|
| 100 | 11.67 | 8.00 | 3.33 | 3.14 | 3.17 |
| 500 | 10.92 | 7.31 | 3.17 | 2.79 | 2.67 |
| 750 | 10.44 | 7.00 | 3.19 | 2.50 | 2.50 |
| 1000 | 11.04 | 7.26 | 3.25 | 2.73 | 2.54 |

Glu

| TM (b0) | P5 | P10 | P15 | P20 | P30 |
|---|---|---|---|---|---|
| 100 | 8.17 | 5.75 | 3.22 | 5.29 | 3.17 |
| 500 | 7.83 | 5.69 | 3.22 | 4.79 | 3.25 |
| 750 | 7.88 | 5.71 | 3.26 | 4.05 | 3.28 |
| 1000 | 7.96 | 5.94 | 3.31 | 4.57 | 3.50 |

| TM (b0+3) | P5 | P10 | P15 | P20 | P30 |
|---|---|---|---|---|---|
| 100 | 9.33 | 7.25 | 3.22 | 5.86 | 4.00 |
| 500 | 8.92 | 6.81 | 3.17 | 5.14 | 3.92 |
| 750 | 9.19 | 6.79 | 3.26 | 4.32 | 4.00 |
| 1000 | 9.65 | 6.87 | 3.33 | 5.23 | 4.08 |

tCr

| TM (b0) | P5 | P10 | P15 | P20 | P30 |
|---|---|---|---|---|---|
| 100 | 3.83 | 2.75 | 1.44 | 1.43 | 1.67 |

Tau

| TM (b0) | P5 | P10 | P15 | P20 | P30 |
|---|---|---|---|---|---|
| 100 | 3.17 | 3.25 | 2.33 | 2.14 | 3.00 |

| TM (b0) | P5 | P10 | P15 | P20 | P30 |
|---|---|---|---|---|---|
| 500 | 3.75 | 2.56 | 1.33 | 1.43 | 1.75 |
| 750 | 3.69 | 2.54 | 1.41 | 1.41 | 1.72 |
| 1000 | 3.96 | 2.74 | 1.58 | 1.53 | 1.79 |

| TM (b0+3) | P5 | P10 | P15 | P20 | P30 |
|---|---|---|---|---|---|
| 100 | 4.83 | 3.25 | 2.00 | 1.71 | 2.17 |
| 500 | 4.58 | 3.06 | 1.83 | 1.71 | 2.08 |
| 750 | 4.44 | 3.08 | 1.85 | 1.64 | 2.06 |
| 1000 | 4.74 | 3.16 | 1.97 | 1.70 | 2.13 |

| TM (b0) | P5 | P10 | P15 | P20 | P30 |
|---|---|---|---|---|---|
| 500 | 2.83 | 3.06 | 2.22 | 2.14 | 3.00 |
| 750 | 2.75 | 2.88 | 2.22 | 2.14 | 2.89 |
| 1000 | 2.74 | 2.94 | 2.28 | 2.13 | 2.92 |

| TM (b0+3) | P5 | P10 | P15 | P20 | P30 |
|---|---|---|---|---|---|
| 100 | 4.00 | 4.13 | 2.67 | 2.57 | 4.50 |
| 500 | 3.50 | 3.88 | 2.44 | 2.50 | 3.83 |
| 750 | 3.31 | 3.58 | 2.41 | 2.41 | 3.72 |
| 1000 | 3.30 | 3.48 | 2.42 | 2.43 | 3.79 |

tCho

| TM (b0) | P5 | P10 | P15 | P20 | P30 |
|---|---|---|---|---|---|
| 100 | 5.50 | 5.38 | 4.00 | 4.14 | 4.33 |
| 500 | 5.58 | 5.06 | 3.78 | 4.14 | 4.50 |
| 750 | 5.50 | 4.92 | 3.93 | 4.09 | 4.61 |
| 1000 | 5.78 | 5.06 | 4.50 | 4.23 | 4.75 |

| TM (b0+3) | P5 | P10 | P15 | P20 | P30 |
|---|---|---|---|---|---|
| 100 | 7.33 | 5.63 | 4.22 | 5.00 | 6.67 |
| 500 | 6.58 | 5.44 | 4.06 | 4.64 | 6.00 |
| 750 | 6.56 | 5.33 | 4.11 | 4.50 | 5.89 |
| 1000 | 6.74 | 5.29 | 4.31 | 4.57 | 5.96 |

Ins

| TM (b0) | P5 | P10 | P15 | P20 | P30 |
|---|---|---|---|---|---|
| 100 | 12.17 | 15.50 | 7.89 | 7.71 | 5.50 |
| 500 | 10.00 | 13.44 | 7.28 | 6.93 | 5.50 |
| 750 | 9.75 | 13.42 | 8.11 | 6.27 | 5.39 |
| 1000 | 9.83 | 14.10 | 8.72 | 6.53 | 5.46 |

| TM (b0+3) | P5 | P10 | P15 | P20 | P30 |
|---|---|---|---|---|---|
| 100 | 12.83 | 17.63 | 8.00 | 8.00 | 7.33 |
| 500 | 11.08 | 15.50 | 7.28 | 7.36 | 6.58 |
| 750 | 10.56 | 14.33 | 7.44 | 6.73 | 6.22 |
| 1000 | 10.87 | 14.90 | 8.36 | 6.83 | 6.38 |

**THALAMUS**

tNAA

| TM (b0) | P5 | P10 | P15 | P20 | P30 |
|---|---|---|---|---|---|
| 100 | 6.63 | 3.22 | 2.11 | 2.00 | 2.00 |
| 500 | 7.07 | 3.11 | 2.06 | 2.00 | 2.00 |
| 750 | 7.87 | 3.15 | 2.07 | 2.00 | 2.00 |
| 1000 | 8.55 | 3.28 | 2.08 | 2.00 | 2.00 |

| TM (b0+3) | P5 | P10 | P15 | P20 | P30 |
|---|---|---|---|---|---|
| 100 | 8.38 | 4.22 | 2.33 | 2.56 | 2.78 |
| 500 | 8.07 | 4.00 | 2.17 | 2.28 | 2.59 |
| 750 | 8.00 | 4.04 | 2.15 | 2.19 | 2.50 |
| 1000 | 8.55 | 4.06 | 2.14 | 2.17 | 2.51 |

Glu

| TM (b0) | P5 | P10 | P15 | P20 | P30 |
|---|---|---|---|---|---|
| 100 | 6.13 | 4.00 | 3.11 | 3.00 | 3.33 |
| 500 | 6.27 | 4.06 | 3.06 | 3.06 | 3.29 |
| 750 | 6.48 | 4.11 | 3.07 | 3.11 | 3.35 |
| 1000 | 6.81 | 4.31 | 3.08 | 3.22 | 3.49 |

| TM (b0+3) | P5 | P10 | P15 | P20 | P30 |
|---|---|---|---|---|---|
| 100 | 7.75 | 5.56 | 3.67 | 4.11 | 4.33 |
| 500 | 7.67 | 5.44 | 3.39 | 4.06 | 4.24 |
| 750 | 7.70 | 5.52 | 3.44 | 4.00 | 4.35 |
| 1000 | 7.97 | 5.64 | 3.53 | 4.06 | 4.51 |

tCr

| TM (b0) | P5 | P10 | P15 | P20 | P30 |
|---|---|---|---|---|---|
| 100 | 3.50 | 2.11 | 2.00 | 2.11 | 2.33 |
| 500 | 3.47 | 2.11 | 2.00 | 2.06 | 2.29 |
| 750 | 3.57 | 2.15 | 2.04 | 2.04 | 2.35 |
| 1000 | 3.74 | 2.31 | 2.08 | 2.08 | 2.49 |

| TM (b0+3) | P5 | P10 | P15 | P20 | P30 |
|---|---|---|---|---|---|
| 100 | 4.00 | 2.78 | 2.11 | 2.89 | 2.89 |
| 500 | 3.93 | 2.61 | 2.06 | 2.50 | 2.88 |
| 750 | 3.96 | 2.70 | 2.07 | 2.44 | 2.88 |
| 1000 | 4.16 | 2.78 | 2.14 | 2.56 | 3.03 |

Tau

| TM (b0) | P5 | P10 | P15 | P20 | P30 |
|---|---|---|---|---|---|
| 100 | 2.38 | 2.00 | 2.11 | 3.67 | 5.00 |
| 500 | 2.47 | 2.00 | 2.06 | 3.56 | 5.00 |
| 750 | 2.48 | 2.00 | 2.07 | 3.44 | 4.96 |
| 1000 | 2.61 | 2.06 | 2.11 | 3.47 | 5.17 |

| TM (b0+3) | P5 | P10 | P15 | P20 | P30 |
|---|---|---|---|---|---|
| 100 | 3.13 | 2.89 | 3.00 | 5.78 | 6.67 |
| 500 | 3.00 | 2.56 | 2.78 | 4.94 | 6.47 |
| 750 | 2.91 | 2.41 | 2.74 | 4.63 | 6.27 |
| 1000 | 2.94 | 2.36 | 2.72 | 4.58 | 6.43 |

tCho

| TM (b0) | P5 | P10 | P15 | P20 | P30 |
|---|---|---|---|---|---|
| 100 | 5.88 | 4.11 | 4.11 | 4.44 | 5.44 |
| 500 | 6.40 | 4.06 | 3.94 | 4.56 | 5.29 |
| 750 | 92.87 | 4.30 | 4.00 | 4.48 | 5.12 |
| 1000 | 135.52 | 4.58 | 4.08 | 4.47 | 5.14 |

| TM (b0+3) | P5 | P10 | P15 | P20 | P30 |
|---|---|---|---|---|---|
| 100 | 9.00 | 5.11 | 4.67 | 6.56 | 6.44 |
| 500 | 7.73 | 4.94 | 4.50 | 5.72 | 6.00 |
| 750 | 7.57 | 5.04 | 4.52 | 5.37 | 5.77 |
| 1000 | 7.81 | 5.11 | 4.58 | 5.42 | 5.91 |

Ins

| TM (b0) | P5 | P10 | P15 | P20 | P30 |
|---|---|---|---|---|---|
| 100 | 11.50 | 13.33 | 16.33 | 10.11 | 7.44 |
| 500 | 11.20 | 13.11 | 14.44 | 9.28 | 6.82 |
| 750 | 11.87 | 12.37 | 13.15 | 8.59 | 6.77 |
| 1000 | 12.39 | 12.31 | 13.28 | 8.58 | 6.91 |

| TM (b0+3) | P5 | P10 | P15 | P20 | P30 |
|---|---|---|---|---|---|
| 100 | 12.38 | 16.00 | 25.89 | 21.89 | 9.00 |
| 500 | 11.67 | 13.72 | 18.39 | 15.06 | 8.00 |
| 750 | 11.22 | 12.96 | 15.96 | 12.52 | 7.65 |
| 1000 | 10.87 | 12.89 | 15.44 | 11.81 | 7.74 |

**Table S2:** estimation of $f_{sphere,LT}$ derived using the ratio of the relative thicknesses of the EGL, IGL, PL ("cell body-like" layers) and ML ("process-like" layer). "with WM" means that the white matter layer was included in the "process-like" layers (in addition to ML). Data come from manual measurements of literature cerebellar figures. References are at the bottom of the Supplementary Information.

**Nakayama 2018**[1]

| | P5 | P8/9 | P13/14 | P20/21 | P60 |
|---|---|---|---|---|---|
| with WM | 0.65 | 0.68 | 0.53 | 0.4 | 0.37 |
| without WM | 0.88 | 0.79 | 0.61 | 0.46 | 0.4 |

**Miterko 2019**[2]

| | P5* | P10 | P15 | P20 | P180 |
|---|---|---|---|---|---|
| with WM | 0.75 | 0.59 | 0.47 | 0.42 | 0.33 |
| without WM | 0.75 | 0.63 | 0.5 | 0.44 | 0.4 |

*WM not identifiable in the region of lobule IV/V used for the measure

**Yamanaka 2004**[3]

| | P5 | P7 | P10 | P15 | P20 |
|---|---|---|---|---|---|
| without WM | 0.82 | 0.77 | 0.69 | 0.58 | 0.53 |

**van der Heijen 2021**[4]

| | P7 | | P14 | | Ad |
|---|---|---|---|---|---|
| without WM | 0.82 | | 0.53 | | 0.51 |

**Araujo 2019**[5]

| | P6 | P12 | P15 | | Ad |
|---|---|---|---|---|---|
| without WM | 0.87 | 0.76 | 0.64 | | 0.53 |

**Kim and Scott 2014**[6]

| | | | P14 |
|---|---|---|---|
| without WM | | | 0.43 |

**Supplementary information to Figure 6**
3D reconstruction from Neuromorph
Cerebellum P6 - P9 = 4 neurons
Cerebellum P10 = 6 neurons
Cerebellum P11 = 1 neuron
Cerebellum P12 = 3 neurons
Cerebellum P13 = 3 neurons
Cerebellum P35 = 4 neurons
Cerebellum P37 = 1 neuron
Cerebellum P43 = 1 neuron
Thalamus P9 = 1 neuron
Thalamus P10 = 2 neuron
Thalamus P11= 7 neurons
Thalamus P26 = 9 neurons
Thalamus P30 = 2 neurons
